# Genomic Revisitation and Reclassification of the Genus *Providencia*

**DOI:** 10.1101/2023.12.13.571484

**Authors:** Xu Dong, Huiqiong Jia, Yuyun Yu, Yanghui Xiang, Ying Zhang

## Abstract

Members of *Providencia*, although typically opportunistic, can cause severe infections in immunocompromised hosts. Recent advances in genome sequencing provide an opportunity for more precise study of this genus. In this study, we first identified and characterized a novel species named *Providencia zhijiangensis* sp. nov. It has ≤88.23% average nucleotide identity (ANI) and ≤31.8% in silico DNA-DNA hybridization (dDDH) values with all known *Providencia* species, which fall significantly below the species-defining thresholds. Interestingly, we found that *Providencia stuartii* and *Providencia thailandensis* actually fall under the same species, evidenced by an ANI of 98.59% and a dDDH value of 90.4%. By fusing ANI with phylogeny, we have reclassified 545 genomes within this genus into 20 species, including seven unnamed taxa (provisionally titled Taxon1-7), which can be further subdivided into 23 lineages. Pangenomic analysis identified 1,550 genus-core genes in *Providencia*, with coenzymes being the predominant category at 10.56%, suggesting significant intermediate metabolism activity. Resistance analysis revealed that most lineages of the genus (82.61%, 19/23) carry a high number of antibiotic resistance genes (ARGs) and display diverse resistance profiles. Notably, the majority of ARGs are located on plasmids, underscoring the significant role of plasmids in the resistance evolution within this genus. Three species or lineages (*P. stuartii*, Taxon 3, and *Providencia hangzhouensis* L12) that possess the highest number of carbapenem resistance genes suggest their potential influence on clinical treatment. These findings underscore the need for continued surveillance and study of this genus, particularly due to their role in harboring antibiotic resistance genes.

## Importance

The *Providencia* genus, known to harbor opportunistic pathogens, has been a subject of interest due to its potential to cause severe infections, particularly in vulnerable individuals. Our research offers groundbreaking insights into this genus, unveiling a novel species, *Providencia zhijiangensis* sp. nov., and highlighting the need for a re-evaluation of existing classifications. Our comprehensive genomic assessment offers a detailed classification of 545 genomes into distinct species and lineages, revealing the rich biodiversity and intricate species diversity within the genus. The substantial presence of antibiotic resistance genes in the *Providencia* genus underscores potential challenges for public health and clinical treatments. Our study highlights the pressing need for increased surveillance and research, enriching our understanding of antibiotic resistance in this realm.

## Introduction

*Providencia*, Gram-negative bacteria from the *Enterobacteriales* order, are widespread across diverse environments like water, soil, and animals (1). These bacteria, as opportunistic pathogens, can cause severe human infections, including urinary tract infections and meningitis (2-5). The genus currently encompasses 14 recognized species: *Providencia stuartii*, *Providencia thailandensis*, *Providencia manganoxydans*, *Providencia burhodogranariea*, *Providencia sneebia*, *Providencia hangzhouensis*, *Providencia rettgeri*, *Providencia huaxiensis*, *Providencia wenzhouensis*, *Providencia alcalifaciens*, *Providencia heimbachae*, *Providencia vermicola*, *Providencia entomophila* and *Providencia rustigianii* (6-16). Among these, *P. stuartii*, *P. rettgeri*, and the newly discovered *P. hangzhouensis* are predominantly implicated in human infections.

While prior studies have focused on one or several specific species within the *Providencia* genus (17-19), a comprehensive taxonomic reassessment and deep genomic insight into the entire genus remain elusive. Particularly concerning are the recurrent misclassifications, as evidenced by the incorrect attribution of strains to *P. rettgeri* that in fact belong to *P. hangzhouensis* (16). Such accurate species categorization is foundational to our grasp of bacterial habitat, epidemiology, pathogenesis, and microbiological traits, with vast repercussions across health, industry, and scientific research (20). Thus, there’s an ever-pressing need for up-to-date, meticulously curated taxonomic classifications. Although 16S rRNA gene sequence analysis is a popular tool for bacterial identification, its precision often falls short for accurate species differentiation (21). Owing to the rapid advancement in sequencing technologies, whole-genome-based analyses have become more accessible, gaining prominence in species identification due to their high resolution (22). Average nucleotide identity (ANI), using cutoff values of ≥96%, and in silico DNA-DNA hybridization (dDDH) with a cutoff of ≥70.0% are commonly utilized for precise species identification (23-25).

Despite several studies (16-18) highlighting the challenges in *Providencia* classification, a comprehensive, genome-based investigation into these ambiguities remains absent. To address this deficiency, our study aims to demystify *Providencia* taxonomy, presenting an updated classification and introducing a novel species, *Providencia zhijiangensis* sp. nov. Leveraging the power of genomics, we established a robust genome-based phylogenetic framework to unravel the intricate diversity within the *Providencia* genus. Augmenting our primary focus, we also delved into the exploration of genus-specific core genes, and investigated the landscape of antibiotic resistance and plasmids, offering insights into the multifaceted nature of this genus.

## Results

### Identification and characterization of novel species *Providencia zhijiangensis*

Strain D4759, preliminarily identified as *Providencia alcalifaciens* using the MALDI-TOF MS technique (Vitek MS system, bioMerieux, France), was isolated from a clinical patient’s bile sample. Due to the rarity of its isolation site (as *P. alcalifaciens* is commonly found in diarrhea infections), we subsequently conducted whole-genome sequencing to ascertain its precise species classification.

The D4759 complete genome comprises a chromosome spanning 3,796,440 bp with a G+C content of 42.91% and two plasmids, pD4759_2_1 and pD4759_2_2, measuring 2,683 bp and 2,428 bp in length, respectively. Utilizing the 16S rRNA gene sequences from other *Providencia* species available in the EzBioCloud database (26) and a subsequently constructed phylogenetic tree (**Fig. 1A**), it was confirmed that strain D4759 aligns within the *Providencia* genus. Specifically, D4759 exhibited pronounced 16S rRNA gene sequence similarity to *P. rustigianii* DSM 4542 (99.74%; AM040489) and *P. alcalifaciens* DSM 30120 (99.73%; ABXW01000071). Given the previously stated insufficient resolution of 16S rRNA phylogenies for *Providencia* species delineation (9, 14, 18), we opted for a whole-genome-based phylogenetic analysis approach, comparing D4759 with type strains from the *Providencia* genus. This phylogenetic analysis further substantiated D4759’s placement within the *Providencia* genus, particularly emphasizing its closest association with *P. alcalifaciens* DSM 30120 (**Fig. 1B**). Furthermore, upon comparing the ANI (79.81-88.28%) and dDDH (20.9-31.80%) values of strain D4759 with those of type strains from the *Providencia* species, we found them to be significantly below the thresholds defining species classification [ANI>96%; dDDH>70%] (**Table 1**).

**Figure 1.**
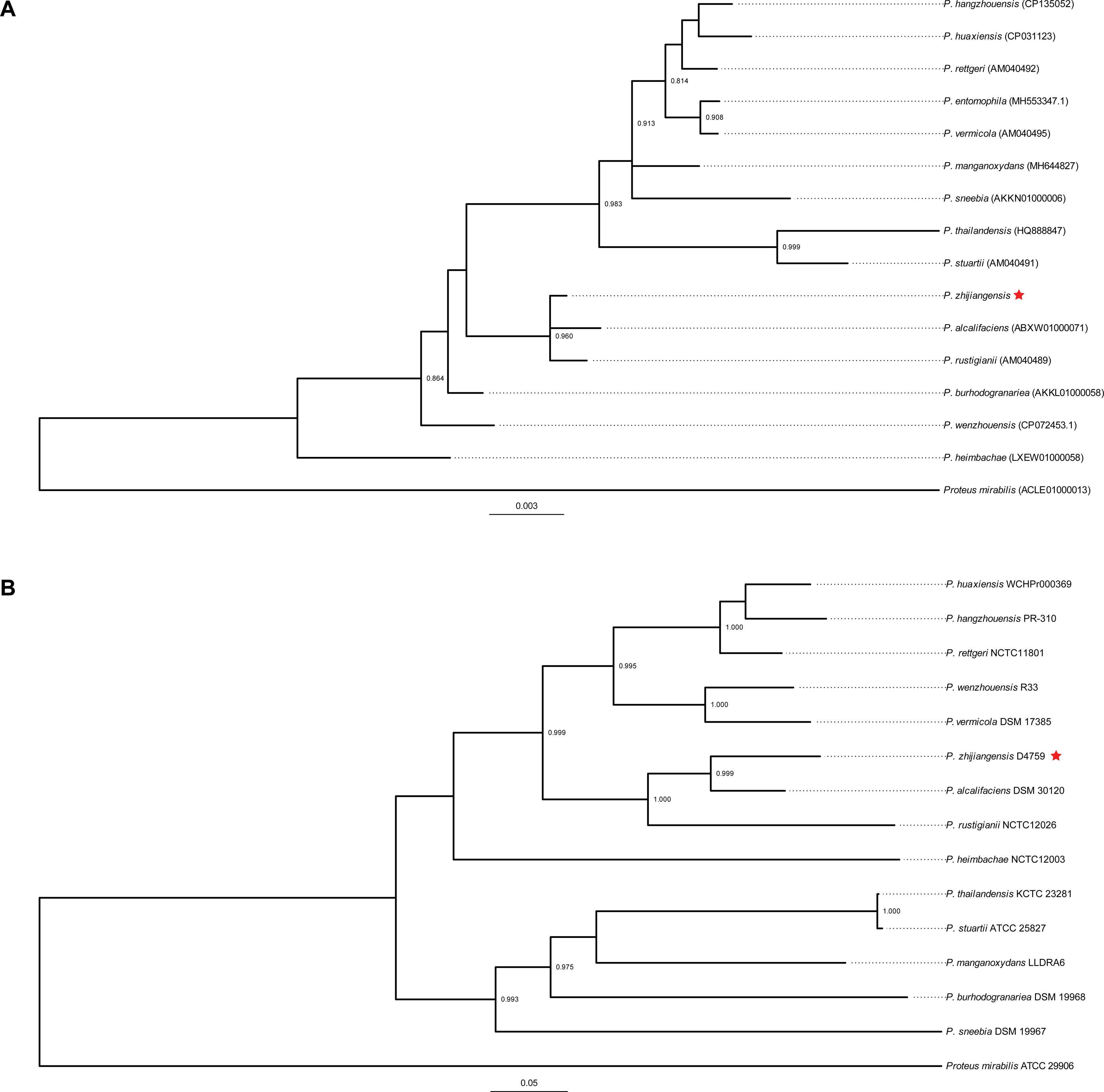
Maximum-likelihood phylogenetic trees of the *Providencia* genus, constructed from (A) 16S rRNA gene sequences and (B) whole-genome sequences. The trees are rooted using *Proteus mirabilis* ATCC 29906 as the outgroup. Branch nodes with bootstrap values >0.8, obtained from 1,000 resamplings, are highlighted. The strain D4759 used in this study is indicated with a red star.

**Table 1.**
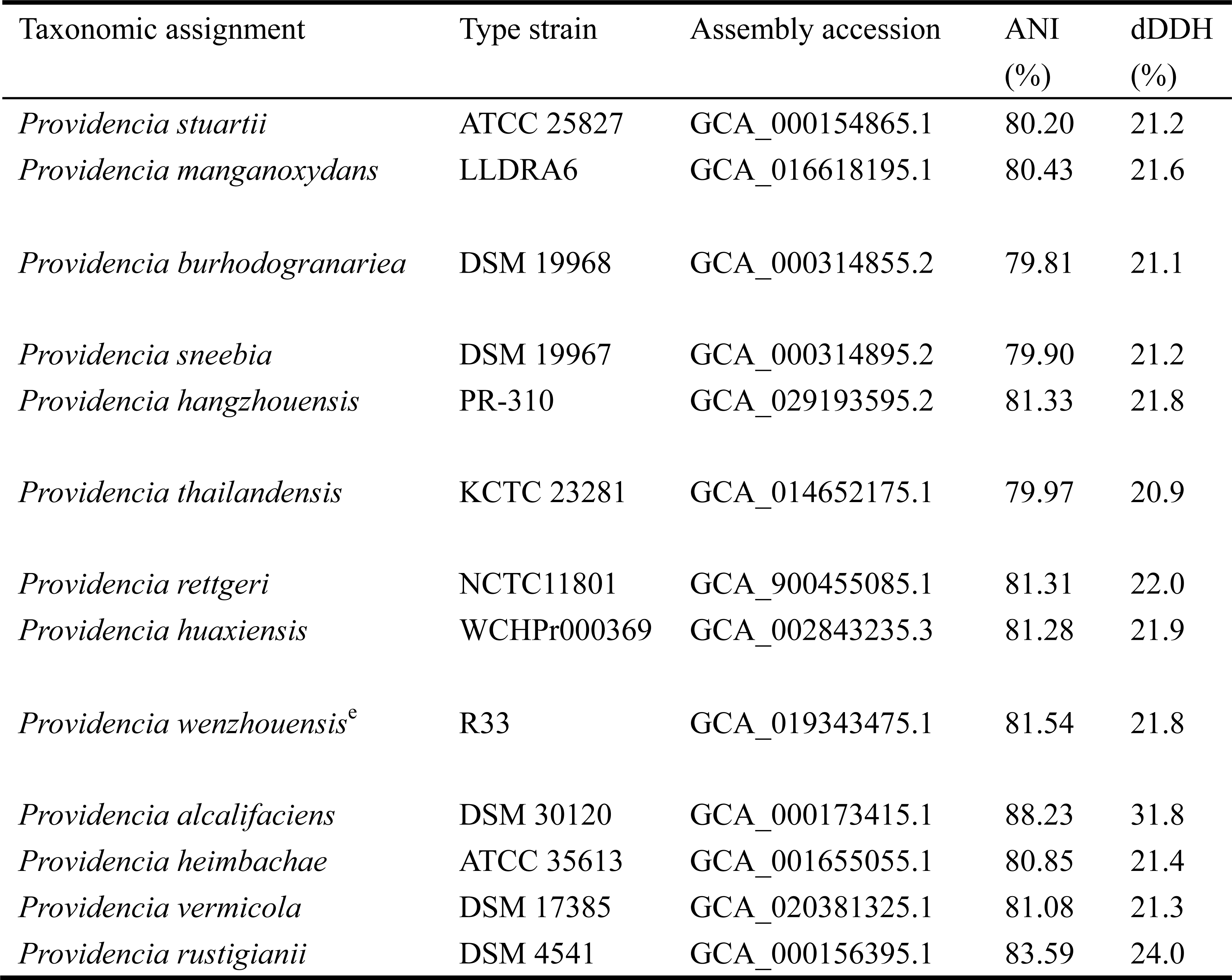
ANI and dDDH values between strains D4759 and the type strains of *Providencia* species.

| Taxonomic assignment | Type strain | Assembly accession | ANI (%) | dDDH (%) |
| --- | --- | --- | --- | --- |
| *Providencia stuartii* | ATCC 25827 | GCA_000154865.1 | 80.20 | 21.2 |
| *Providencia manganoxydans* | LLDRA6 | GCA_016618195.1 | 80.43 | 21.6 |
| *Providencia burhodogranariea* | DSM 19968 | GCA_000314855.2 | 79.81 | 21.1 |
| *Providencia sneebia* | DSM 19967 | GCA_000314895.2 | 79.90 | 21.2 |
| *Providencia hangzhouensis* | PR-310 | GCA_029193595.2 | 81.33 | 21.8 |
| *Providencia thailandensis* | KCTC 23281 | GCA_014652175.1 | 79.97 | 20.9 |
| *Providencia rettgeri* | NCTC11801 | GCA_900455085.1 | 81.31 | 22.0 |
| *Providencia huaxiensis* | WCHPr000369 | GCA_002843235.3 | 81.28 | 21.9 |
| *Providencia wenzhouensis*<sup>e</sup> | R33 | GCA_019343475.1 | 81.54 | 21.8 |
| *Providencia alcalifaciens* | DSM 30120 | GCA_000173415.1 | 88.23 | 31.8 |
| *Providencia heimbachae* | ATCC 35613 | GCA_001655055.1 | 80.85 | 21.4 |
| *Providencia vermicola* | DSM 17385 | GCA_020381325.1 | 81.08 | 21.3 |
| *Providencia rustigianii* | DSM 4541 | GCA_000156395.1 | 83.59 | 24.0 |

The biochemical characteristics of strain D4759 and all other known *Providencia* species are detailed in **Table 2**. D4759 grows between 15 to 42°C, exhibiting optimal growth at 35 and 37°C. Morphologically, it consists of Gram-negative, motile, facultatively anaerobic rods. On nutrient agar incubated for 24 hours at 37°C, colonies appeared circular, raised, yellow, opaque, and with a smooth texture. Notably, D4759 lacks oxidase activity. D4759 can utilize acid from D-glucose, sucrose, and glycerol, but not from D-mannitol, inositol, D-sorbitol, L-rhamnose, melibiose, amygdalin, D-arabinose, aesculin, arbutin, cellobiose, 2-ketogluconate, D-Lyxose, salicin, or D-xylose. A positive reaction is observed for deaminase, while negative reactions are found for β-galactosidase, arginine dihydrolase, ornithine decarboxylase, lysine decarboxylase, and gelatinase. The Voges-Proskauer reaction for D4759 is positive, but it shows negative results for urease activity and indole production. D4759 can utilize citrate but does not produce H_2_S. D4759 exhibits distinct biochemical characteristics compared to other *Providencia* species. The strain D4759 stands out due to its positive response to acetoin production tests and its lack of indole production. When contrasting D4759 with clinically more prevalent *Providencia* species, specifically *P. stuartii*, *P. rettgeri*, and *P. hangzhouensis*, it is noteworthy that D4759 does not utilize inositol and indole. This distinctive biochemical profile could serve as a rapid diagnostic criterion to differentiate D4759 from its close relatives in clinical settings. The antimicrobial resistance susceptibility tests shows that D4759 is sensitive to all tested antibiotics except tigecycline (**Table S1**). The total fatty acid profile shows that there are two predominant fatty acids (>10 %), i.e., C_16:0_ (35.03%) and C_14:0_ (11.41%).

**Table 2.**
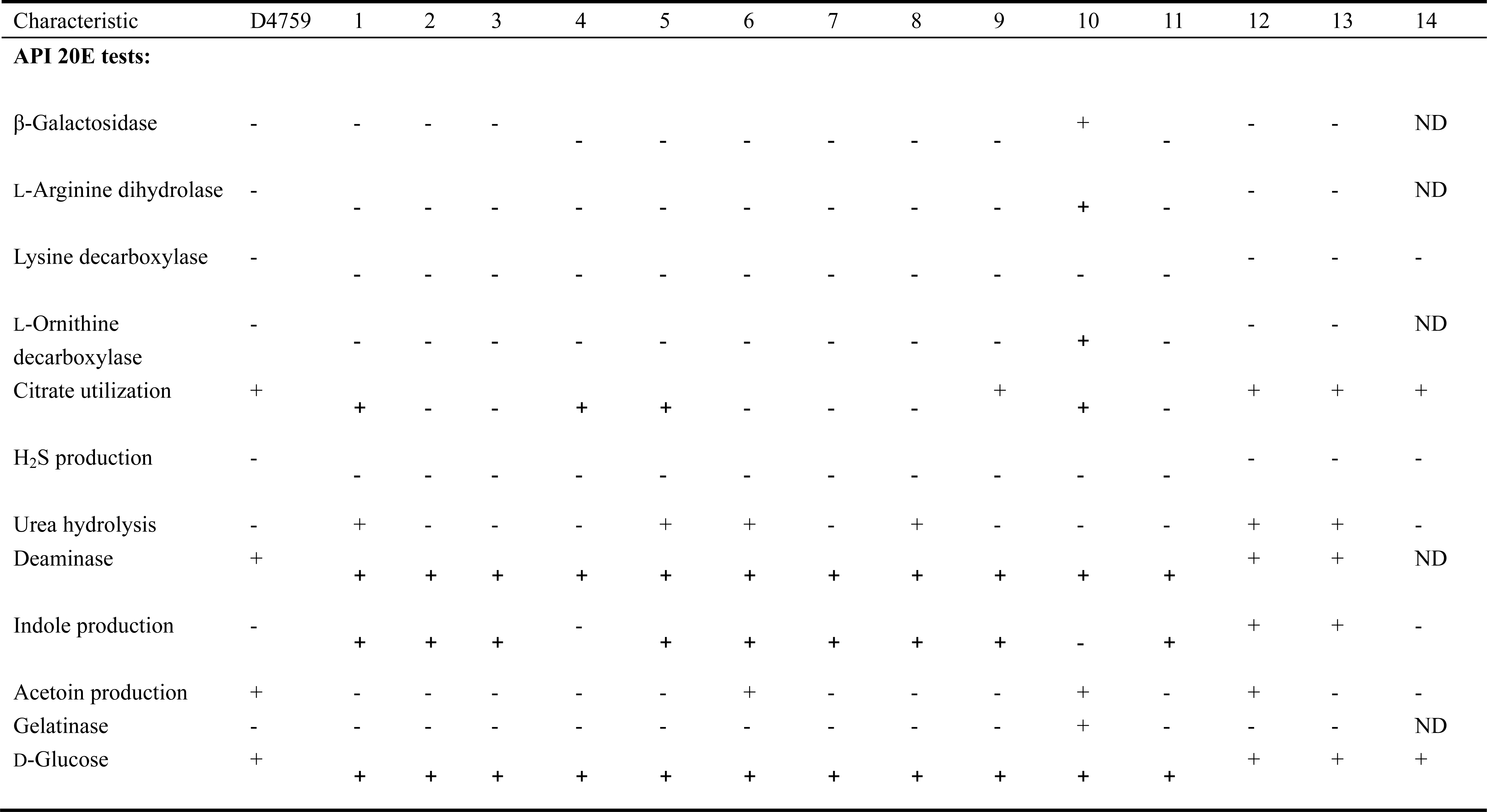
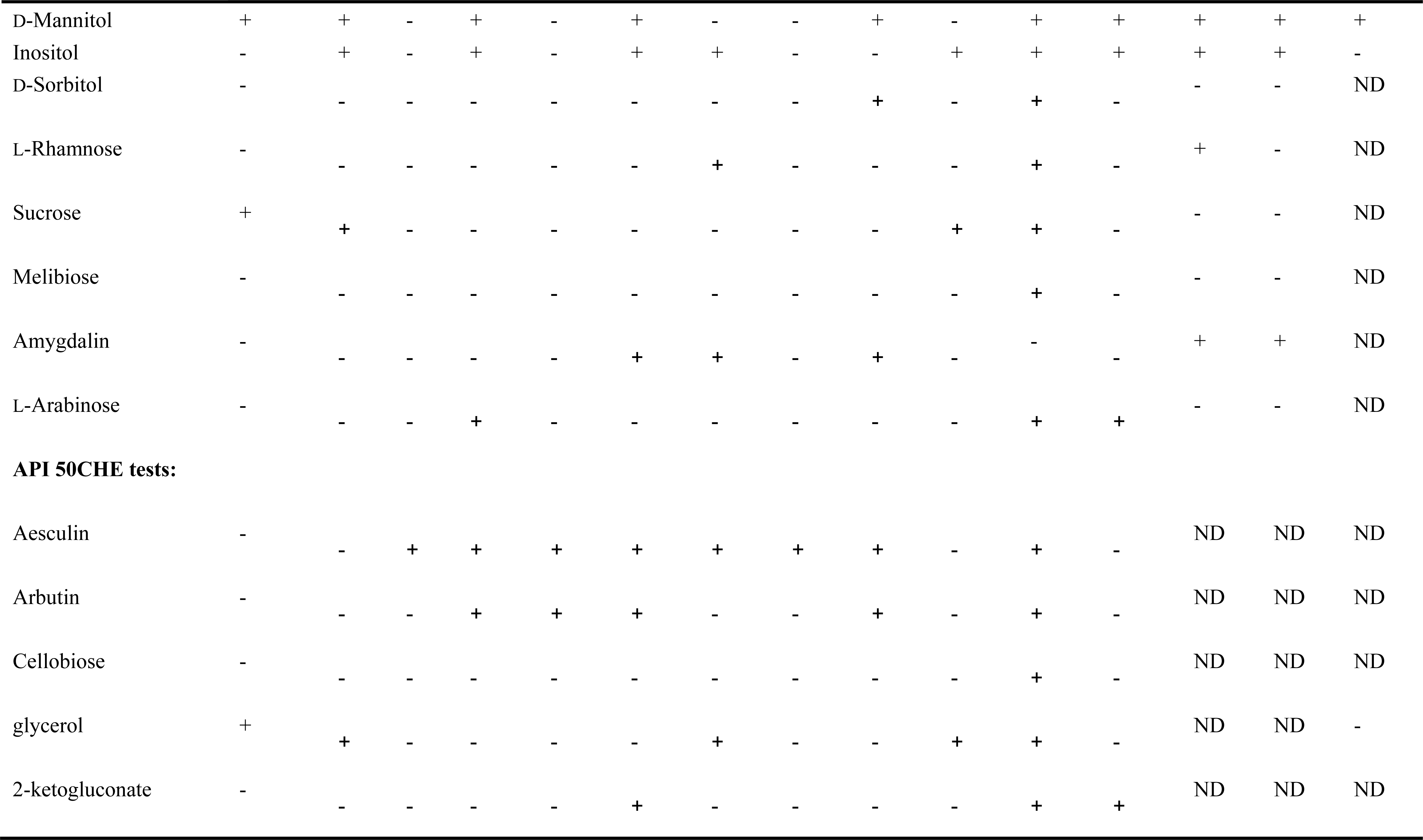
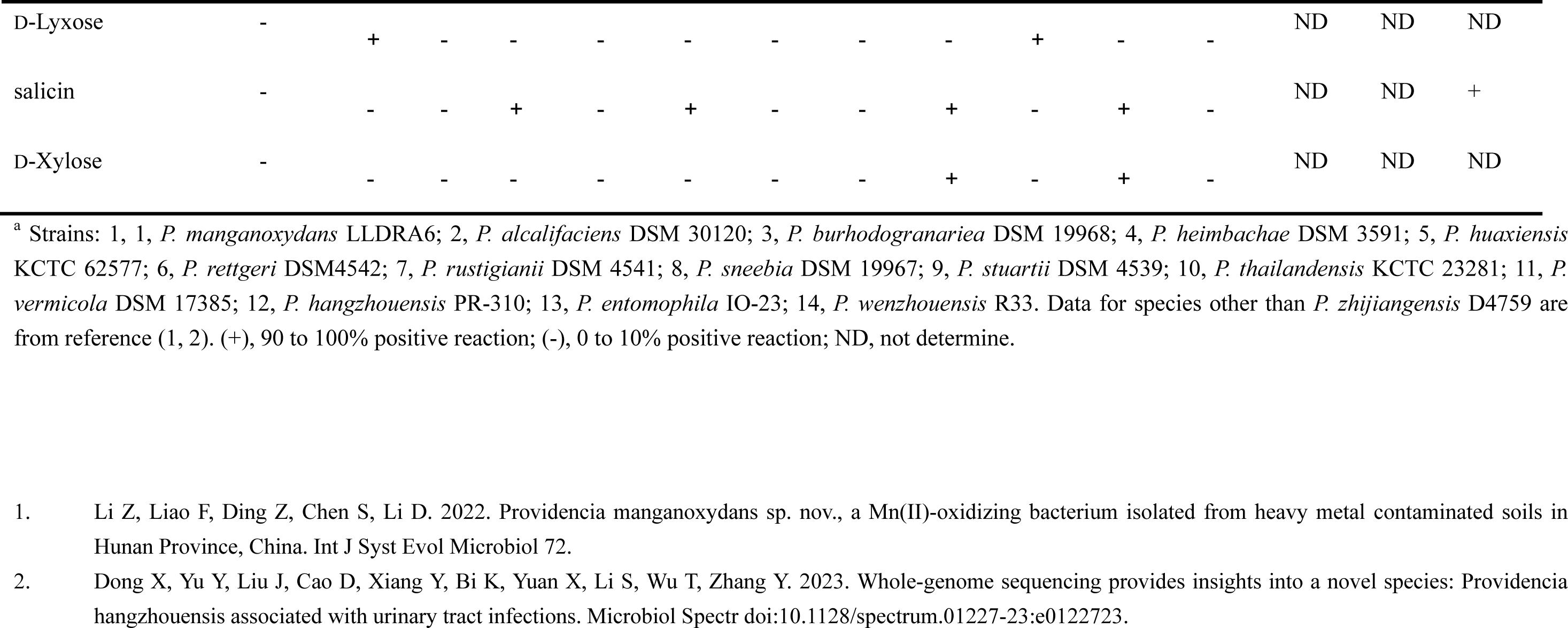
Biochemical characteristics of strain D4759 and type strains of other *Providencia* species^a^.

Based on both genotypic and phenotypic characterizations, we propose that strain D4759 be recognized as a novel species, for which we suggest the name *Providencia zhijiangensis* sp. nov. (zhi.jiang.en’sis. N.L. masc. adj. zhijiangensis, referring to the Xihu District of Hangzhou City, Zhejiang Province, China).

### Curation of *Providencia* genomes with the updated taxonomy

Following the novel species identification and considering the noted inaccuracies in previous classifications (16), a re-evaluation is paramount. From our comprehensive collection approach, we sequenced other three recently collected *Providencia* clinical strains (two from urine and one from tissue fluid). Additionally, after quality assessment, 541 out of the 735 *Providencia* genomes from GenBank were included for a thorough species identification analysis.

By comparing the pairwise ANI of the genomes, we categorized all assemblies into 20 phylogroups using a threshold of 96%. The pairwise dDDH between each phylogroup was found to be less than 70%. Interestingly, when we further determined the species name of each phylogroup by Type Strain Genome Server (TYGS), we found that phylogroup 1 corresponded to both *Providencia stuartii* (dDDH: 94.70%) and *Providencia thailandensis* (dDDH: 94.20%). A comparison of type strains of *P. thailandensis* [KCTC 23281 (GCA_014652175.1)] and *P. stuartii* [ATCC 25827 (GCA_000154865.1)] revealed that their dDDH and ANI values were 94.10% and 99.11%, respectively, suggesting that they indeed constitute a single species. Following the principles outlined by the International Code of Nomenclature of Bacteria (ICNP) (27), *P. stuartii* (6) holds priority over *P. thailandensis* (7) for the species name. Consequently, we propose that *P. thailandensis* should be considered a later heterotypic synonym of *P. stuartii*. These results suggested the taxonomy of the *Providencia* genus could be updated to comprise 20 species, encompassing seven unnamed taxa (Taxon 1-7; **Table 3**).

**Table 3.** Updated genomic species affiliated to genus *Providencia^a^*.

| phylo group | Taxonomic assignment<sup>b</sup> | Type strain<sup>c</sup> | G + C mol%<sup>d</sup> | Assembly accession | No. of genomes |
| --- | --- | --- | --- | --- | --- |
| 1 | *Providencia stuartii* | ATCC 25827 | 41.47 | GCA_000154865.1 | 87 |
| 2 | **Taxon 7** | PROV260 | 40.92 | GCA_028477965.1 | 25 |
| 3 | *Providencia manganoxydans* | LLDRA6 | 40.12 | GCA_016618195.1 | 6 |
| 4 | *Providencia burhodogranariae* | DSM 19968 | 39.08 | GCA_000314855.2 | 1 |
| 5 | **Taxon 1** | BIGb0506 | 39.86 | GCA_025961275.1 | 1 |
| 6 | *Providencia sneebia* | DSM 19967 | 38.06 | GCA_000314895.2 | 1 |
| 7 | *Providencia hangzhouensis* | PR-310 | 40.43 | GCA_029193595.2 | 137 |
| 8 | ***Providencia zhijiangensis*** | D4759 | 42.85 | GCA_030315915.1 | 5 |
| 9 | **Taxon 2** | PR_162 | 40.92 | GCA_003936755.2 | 4 |
| 10 | *Providencia rettgeri* | NCTC11801 | 40.35 | GCA_900455085.1 | 103 |
| 11 | **Taxon 3** | PROV167 | 40.27 | GCA_028480775.1 | 80 |
| 12 | *Providencia huaxiensis* | WCHPr000369 | 40.83 | GCA_002843235.3 | 16 |
| 13 | *Providencia wenzhouensis*<sup>e</sup> | R33 | 41.47 | GCA_019343475.1 | 6 |
| 14 | **Taxon 4** | PROV227 | 41.32 | GCA_028479185.1 | 9 |
| 15 | *Providencia alcalifaciens* | DSM 30120 | 41.86 | GCA_000173415.1 | 31 |
| 16 | *Providencia heimbachae* | ATCC 35613 | 40.41 | GCA_001655055.1 | 5 |
| 17 | *Providencia vermicola* | DSM 17385 | 41.17 | GCA_020381325.1 | 2 |
| 18 | *Providencia rustigianii* | DSM 4541 | 41.29 | GCA_000156395.1 | 9 |
| 19 | **Taxon 5** | JGM172 | 42.55 | GCA_018257405.1 | 16 |
| 20 | **Taxon 6** | 2019-04-29291-1-1 | 42.55 | GCA_915403165.1 | 1 |

a: Owing to the unavailability of a genome for *Providencia entomophila*, this species was excluded from the updated classification.
b: Taxon identified in this study is marked in bold.
c: The type strain is selected based on the earliest time of release or the deposited type strain in the NCBI GenBank database.
d: Average GC content within species
e: This species name is not valid enough.

The majority strains of *Providencia* (n = 137, 25.14%) were identified as *P. hangzhouensis*. This was followed by *P. rettgeri* (n = 103, 18.90%), and *P. stuartii* (n = 87, 15.96%; **Table 3**). Notably, a substantial portion of genomes (n = 136, 24.95%) could not be classified under existing species names. We have tentatively named these as Taxon 1-7, with Taxon 3 being the most prevalent (n = 80, 58.82%; **Table 3**). Further characterization of Taxon 1 through 7 using phenotypic methods is necessary to establish their species status and assign proper species names, in accordance with the current ICNP (27).

However, the increase in newly recognized species presents a new question: does this genus stand as a separate entity, or is it a conglomerate of multiple genera? To further scrutinize the *Providencia* genus, we calculated the pairwise average amino acid identity (AAI) values among all genomes within this genus. The values were found to be greater than or equal to 81.26%, surpassing the acknowledged genus-level threshold, which typically ranges between 65% and 72% AAI (28). This means that *Providencia* presently is indeed a distinct genus comprising 20 species.

### Genome diversity within the *Providencia* genus

Variations in genome size, protein-coding sequences, and GC content both within and between species indicate the presence of significant genetic diversity within the *Providencia* genus (**Table 3, Table S2, Fig. S1**). To gain insight into the genome diversity of the *Providencia* genus, a maximum likelihood (ML) phylogeny was constructed from a concatenated core-gene alignment comprised of 2077 loci present (covering 2.23M nucleotides) in 95% of the 545 *Providencia* strains collected in our study. Deep divisions within this phylogeny, as depicted in **Figure 2**, correspond to both the updated genus taxonomy described above and species-level grouping calculated using ANI with a cutoff of 96% for clustering. Applying a Hierarchical Bayesian Analysis of Population Structure (BAPS) to the core single-nucleotide polymorphism (SNP) results, we clustered the 545 genomes into 23 monophyletic lineages, designated L1-L23.

**Figure 2.**
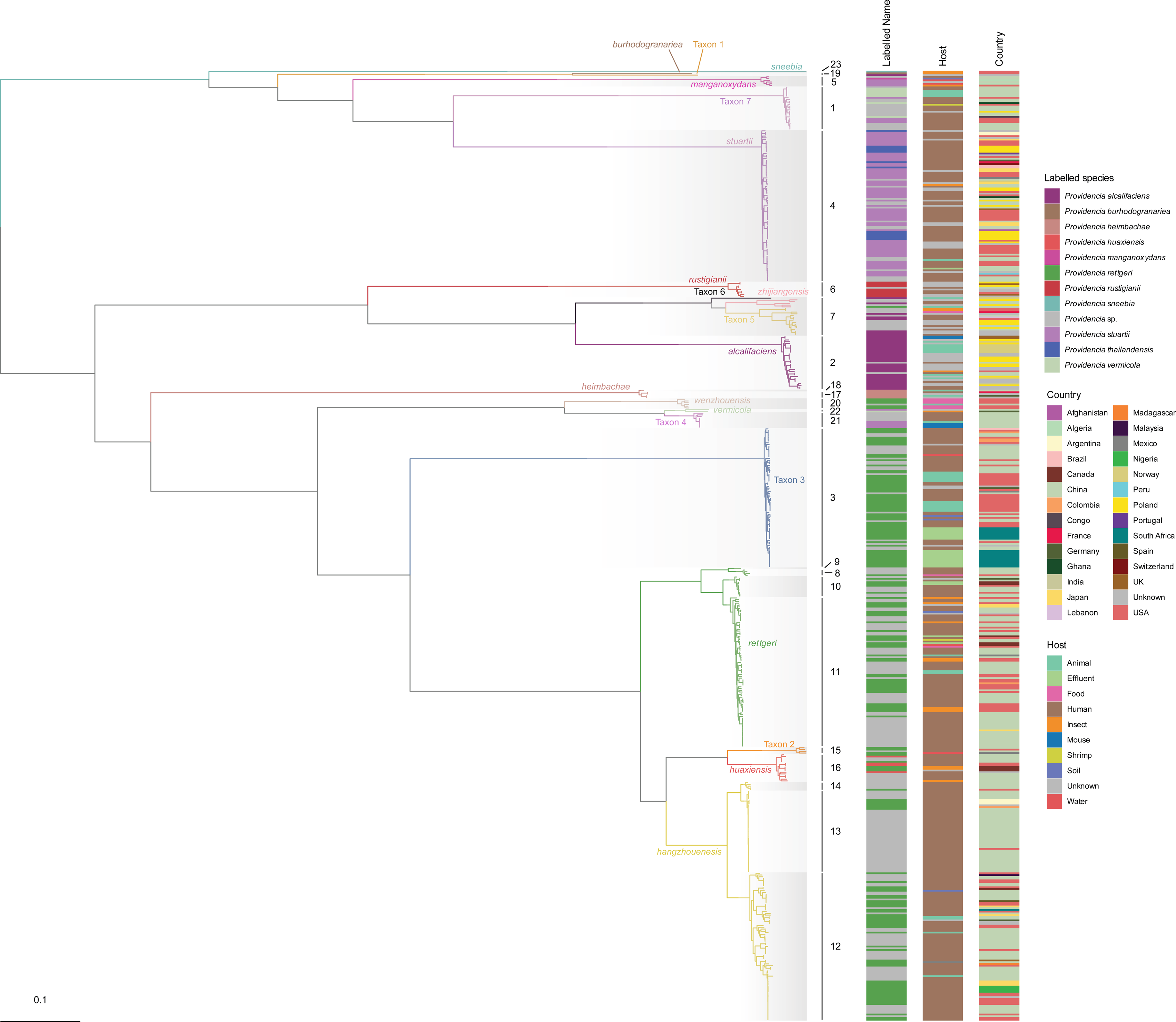
The ML phylogeny of the *Providencia* genus based on the core genome alignment of 545 isolates. The branch colors correspond to phylogroups, which are clustered by ANI, while the shading of clades represents lineages as divided by fastbaps. Unless otherwise specified, the color schemes for all species and lineages in this study are in accordance with those established in Fig. 2.

Intriguingly, the phylogenetic analysis revealed that L19 are composed of three species, including *P. zhijiangensis* and Taxon 5-6 (**Fig. 2**). A comparison of pairwise ANI within these species has shown ANI values ranging between 94.25 and 94.63%. These values approached the threshold typically used to delineate species boundaries, implying a close relationship among these species.

### Pangenome analysis of *Providencia* genus

Prior pan-genomic studies on individual *Providencia* species have highlighted significant variability within the pan-genome (17). After a comprehensive reclassification of the entire Providencia genus, we constructed its pan-genome using a population structure-aware approach (29). The pangenome of the 545 *Providencia* strains comprises 34,087 gene families, with 2,077 of these designated as traditional core genes due to their presence in at least 99% of all genomes. However, when considering only those core genes present in more than 95% of isolates within each lineage, a total of 1,550 genes emerged as core to all lineages, thereby forming the collection core (**Fig. 3A, B**). Collectively, these 1,550 core genes represent approximately a third of genes in an individual isolate and constitute 4.55% of the total pangenome. The pan-genome accumulation curves for species with multiple instances showed an open pan-genome tendency in almost all species, excluding Taxon 4 (**Fig. 3C**). Our pan-genome analysis also identified species or lineage-specific genes (**Fig. 3B**). For instance, in *P. stuartii* and Taxon 3, a distinct set of unique core genes was observed, contrasting with other species (**Fig. S2, S3**). Conversely, a significant proportion of genes in Taxon 7 are also found in other species, primarily in *P. stuartii* (**Fig. S4**). Additionally, in *P. hangzhouensis* L13 and L14, most of the core lineage genes are spread across various lineages (**Fig. S5, S6**).

**Figure 3.**
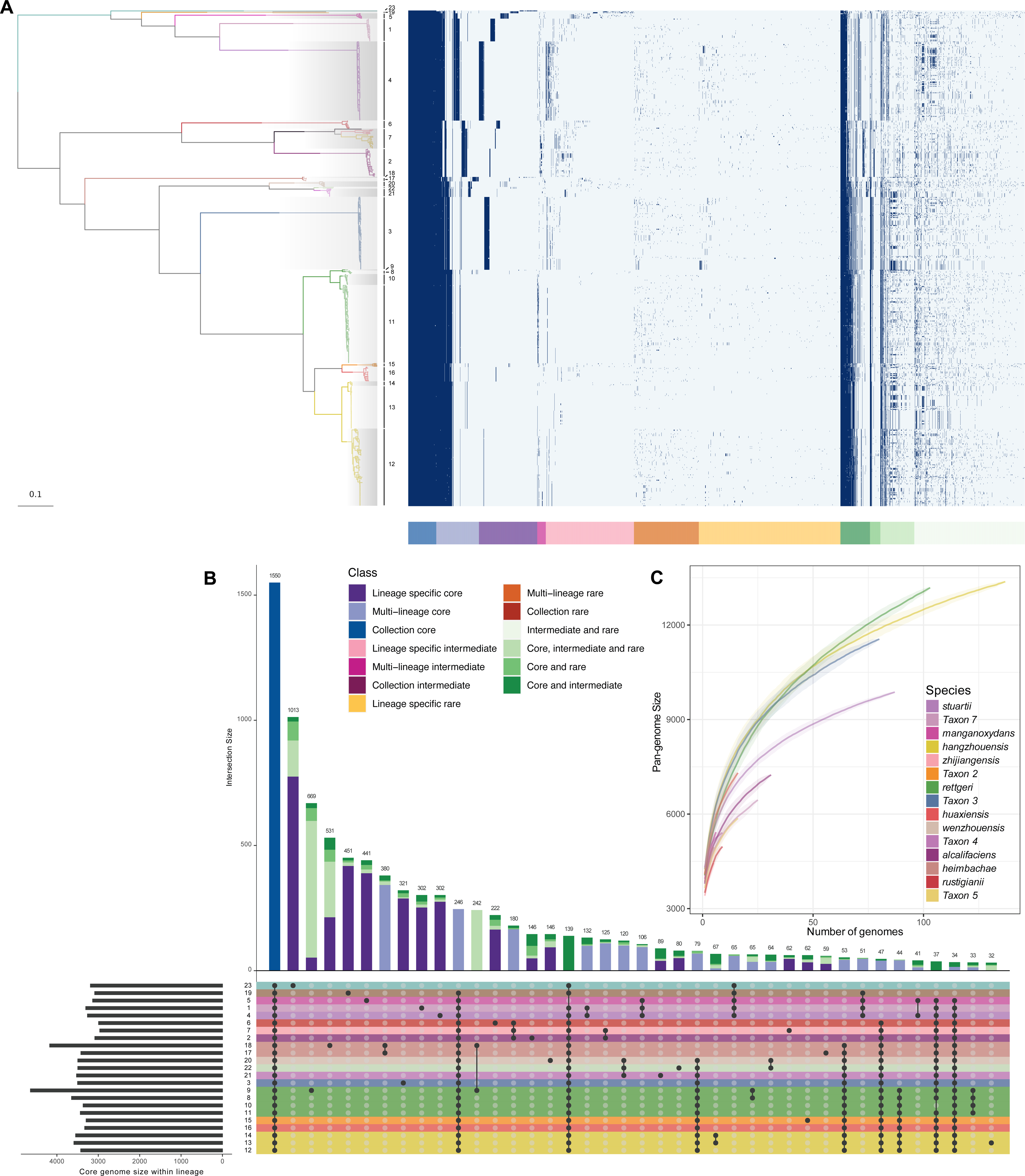
The pangenome analysis of *Providencia* genus. **(A)** Pangenome gene presence and absence heatmap along with the ML tree. The arrangement of genes in the heatmap adheres to the gene classification as defined by twilight (29). **(B)** A UpSetR plot of Intersections of lineage-specific core genomes generated by UpSetR package (65). Each row symbolizes a distinct lineage, while each column indicates their intersections within the matrix, denoted by black dots. **(C)** Pangenome accumulation curves of each species (n>1).

### Functional characterization of the *Providencia* genus-core genes

Functional analysis of the 1,550 core genes of the *Providencia* genus was conducted using the Clusters of Orthologous Groups (COG) approach, aiming to shed light on the foundational functional characteristics and metabolic abilities of the genus. The analysis revealed that these core genes predominantly support essential cellular processes, encompassing coenzyme metabolism (30), inorganic ion transport, and DNA replication (**Fig. 4A**). Among these, coenzymes stood out as they made up the most substantial proportion at 10.56%, emphasizing a pronounced activity of the *Providencia* genus in intermediate metabolism. An additional 4.82% of genes associated with “replication, recombination and repair” could be indicative of potential horizontal gene transfer (HGT) events (31). Notably, a considerable fraction, 15.44%, of the core genes has yet to be assigned specific functions, suggesting uncharted territories in the functional landscape of the genus that beckon further exploration.

**Figure 4.**
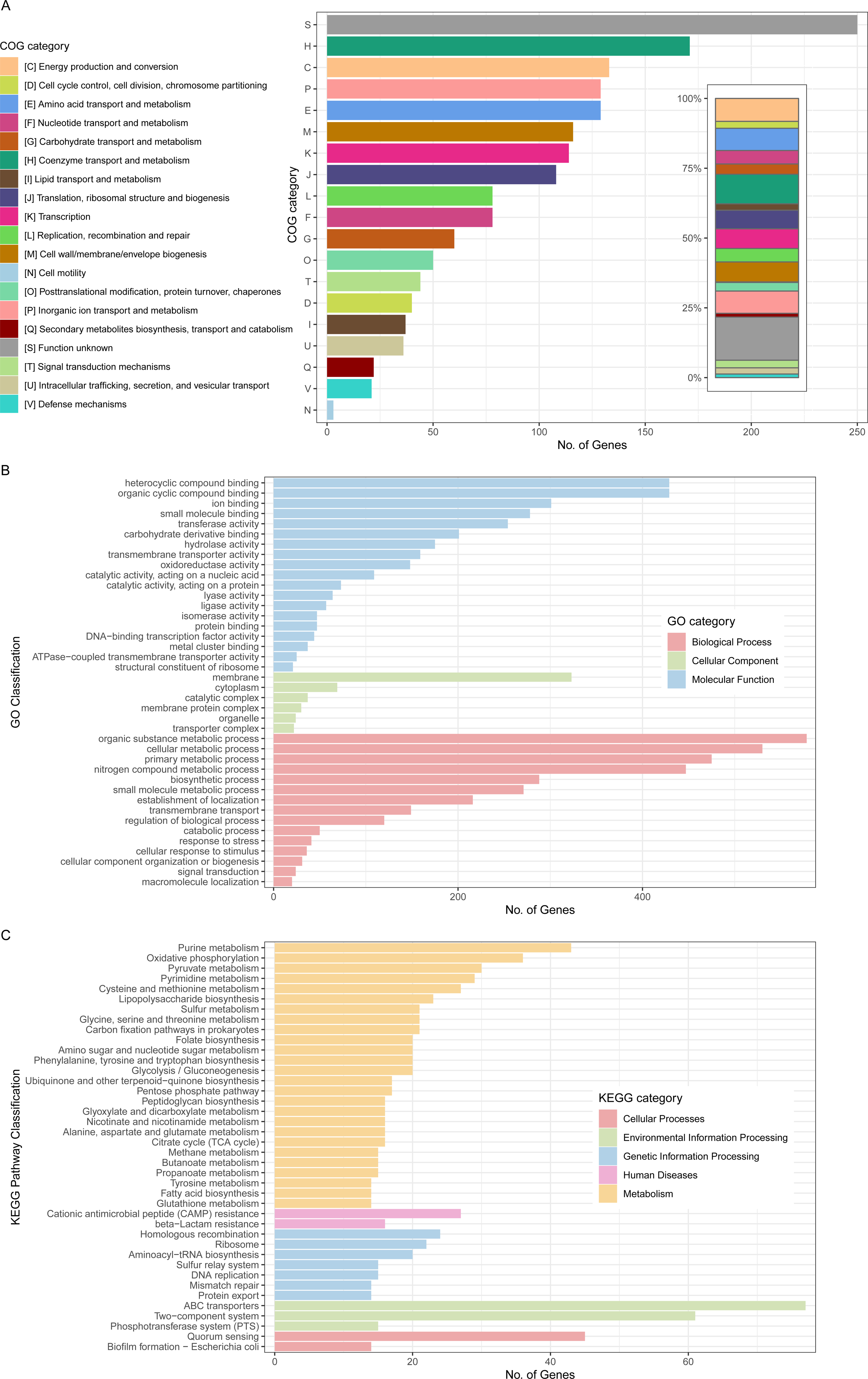
The bar chart presents the functional annotation of 1550 core genes, systematically arranged following the COG **(A)**, GO **(B)**, and KEGG **(C)** classification schemes.

Supplementary insights from Gene Ontology (GO) and Kyoto Encyclopedia of Genes and Genomes (KEGG) analyses complemented our understanding. There was a marked enrichment in metabolic processes, unveiling key metabolic pathways in the genus, spanning organic substance, cellular, and nitrogen compound metabolism (**Fig. 4B**). Interestingly, the KEGG analysis also brought to light pathways related to ‘beta-Lactam resistance’ (**Fig. 4C**), suggesting a potential tendency within the genus towards resistance against beta-lactams, a critical class of antibiotics.

### Resistome characteristics of *Providencia* genus

Antibiotic resistance has been documented in various degrees among *Providencia* species in clinical settings (32, 33). In an effort to understand the landscape of antibiotic resistance genes (ARGs) in the *Providencia* genomes, we performed an analysis on 545 strains, primarily sourced from public databases. Of these, we identified 164 genes spanning 14 categories of antibiotic resistance, ranging from aminoglycosides to tigecycline (**Fig. S7**). Overall, the presence of ARGs was detected in 399 genomes (73.21%), encompassing the vast majority of lineages, with the exception of L6 (*P. rustigianii*), L19 (*P. burhodogranariea* and Taxon 1), L20 (*P. wenzhouensis*) and L23 (*P. sneebia*). In our attempt to find distinctive resistance genes unique to each species or lineage, no such specific genes were discovered. Despite this, *P. stuartii* L4 presented an interesting scenario. We noticed a high frequency of three specific genes—*aac(2’)-Ia*, *tet(B)*, and *catA3*—in this lineage. More precisely, *aac(2’)-Ia* was found in 98.85% of the *P. stuartii* genomes, *tet(B)* in 96.55%, and *catA3* in 94.25%. The prevalent presence of these genes in this lineage may provide a plausible explanation for the frequent categorization of *P. stuartii* as a multidrug-resistant organism.

Beta-lactams, commonly employed in clinical practice, are often central to the treatment of resistant infections, with carbapenems being the last resort. In light of this, we focused our study on carbapenem-resistance genes and identified 17 such genes. Among these, *bla*_IMP-27_ and *bla*_NDM-1_ were the most prevalent (**Fig. S7, S8**). Our investigation into the diversity and abundance of these genes revealed that the clinical lineages L4 (*P. stuartii*) and L12 (*P. hangzhouensis*), primarily of human origin (>75%), exhibited a higher diversity of genes (47.06%, 8/17) compared to other lineages. This finding suggests that these lineages possess greater number of antibiotic resistance genes, warranting further investigation. Additionally, despite the non-clinical lineage L3 (Taxon 3) having only 50% human origin, we identified a significant presence of carbapenem resistance genes in it (**Fig. S8, S9**). The preponderance of these resistance genes in this lineage underscores the need for heightened vigilance regarding the emergence of this species in the clinical setting.

### Contribution of plasmids to the *Providencia* genus

To elucidate the potential impact of plasmids on the *Providencia* genus, we conducted a meticulous examination of plasmid contigs within 545 genomes. Our investigation led to the identification of 721 putative plasmids across 342 genomes spanning 16 distinct species, of which 294 (40.78%) are present in *P. hangzhouensis*. The plasmid dataset displays a broad spectrum of sizes (∼1–240 kb) and GC content (∼28–62%), highlighting their diversity within the *Providencia* species (**Supplementary Data 2**, **Fig. 5B, S10B-C**).

**Figure 5.**
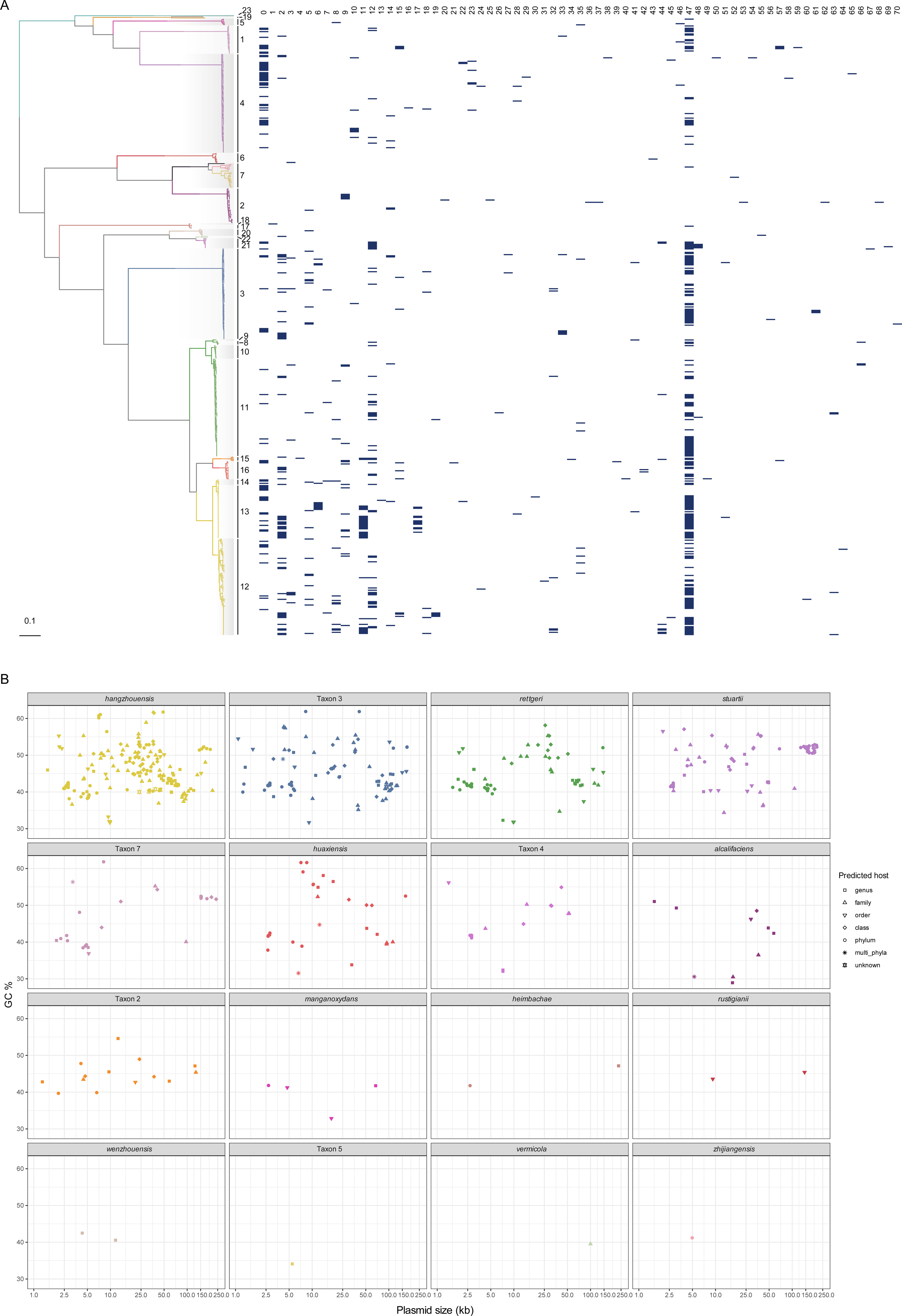
Predicted plasmids across *Providencia* genus. **(A)** Distribution of the 71 plasmid clusters against the ML phylogeny. **(B)** Distribution map of plasmids carried by various species. Plasmid size is denoted on the horizontal axis, whereas the vertical axis corresponds to the GC content. Distinct shapes are employed to symbolize various predicted host sources.

Cluster analysis of the plasmids revealed that the 721 plasmids could be further categorized into 71 clusters (**Fig. 5A**). Intriguingly, cluster 47, identified as the Col3M plasmid, emerged as the most dominant, comprising 29.40% (212/721) of the total, and was found in a major fraction of species harboring plasmids (68.75%, 11/16). Further investigation into the average GC content of the ColM plasmid revealed a similarity with the average GC content of its genus, suggesting that this plasmid may be a genus-specific carrier (34, 35). Though only 24.41% of the identified plasmids were predicted to be either conjugative or mobilizable, their hosts were diverse, spanning across multiple genera and phyla (**Fig. S10A**). This observation could be attributable, in part, to an insufficient exploration of this genus’s plasmids and potentially to frequent HGT activity within the plasmid, leading to loss of the corresponding transfer element.

Given the significance of plasmids in the dissemination of ARGs, we focused our attention on the distribution of ARGs within these clusters. Of the 3757 ARGs, 2019 (54.27%) were identified on plasmids, indicating a heightened burden of ARGs carried by these entities. A standout observation centered on cluster 0. Not only did this cluster claim the highest count of resistance genes, encompassing an impressive 36.97% of the overall ARGs, but it also emerged as a primary reservoir for carbapenemase genes, with a notable 34 out of the 66 identified carbapenemase genes localized within it (**Fig. S11**). Intriguingly, cluster 0 predominantly consists of large MOBH conjugative plasmids that are directly associated with the globally disseminated MDR-associated IncC type plasmids (36), further emphasizing the significant clinical implications of our findings.

Furthermore, we found that plasmids bore a higher load (43.79%, 645/1473) of insertion sequences (ISs) associated with HGT. Certain ISs demonstrated a strong association with plasmids, with the most robust correlation observed for IS*26* (**Fig. S12**). Previous research has established that IS*26* can instigate plasmid reorganization in clinical settings via replicative transposition (37), as well as amplify ARGs (38).

## Discussion

In this study, we first identified and characterized a novel species, *Providencia zhijiangensis* sp. nov. Intriguingly, upon combining phylogeny and overall genome relatedness indexes (OGRIs) (including ANI and dDDH), we found that *P. stuartii* and *P. thailandensis* actually belong to the same species, contradicting the results of the biochemical identification conducted by Khunthongpan et al. (7). This discrepancy could be attributed to variations in measurement conditions or intra-species differences. Through our analysis of ORIGs and phylogeny, we unearthed a remarkable finding: the *Providencia* genus, previously assumed to comprise only 14 species, in fact includes 20 species. This updated count includes the our previously identified *P. hangzhouensis* (39), along with seven unnamed taxa (Taxon 1-7) which warrant further phenotype-based characterizations. Additionally, upon precise identification, we found that the labels of a large number of genomes are inconsistent with their actual species (**Fig. 2**), indicating significant misclassification. This is especially evident in the case of *P. rettgeri*, where 166 strains initially labeled as *P. rettgeri* were in fact primarily composed of the species *P. hangzhouensis* and Taxon 3 (**Fig. 2**, **Supplementary Data 1**). These observations underscore the prevailing confusion in the current classification of *Providencia*, highlighting the importance of accurate reclassification of species within this genus as accomplished in this study.

The considerable variance in genomic characteristics among different species and lineage divisions suggest a diverse genome within the genus. It appears unusual that lineage 19 occupies the ecological niche of *P. zhijiangensis* and Taxon 5-6, although ANI values approaching the species threshold (94.25-94.63%) indicate a high degree of similarity between these three species. Nonetheless, further internal classification requires a larger pool of genomes.

The significant diversity observed in the *Providencia* pan-genome, as underscored by earlier research (17), suggests notable genomic plasticity within the *Providencia* species. This plasticity is further reflected in our findings, where a significant portion of the pan-genome remains open. It is intriguing that, despite sharing a phylogenetic branch and a similar core gene count with *P. wenzhouensis*, Taxon 4 exhibits a closed pangenome, whereas *P. wenzhouensis* has an open one. This could shed light on distinct evolutionary pathways or pressures faced by these lineages.

Species-specific genes frequently delineate the unique ecological niche of individual organisms, a notion reflected in our pan-genome analysis which uncovers species or lineage-specific genes (**Fig. 3B**). Our analysis reaffirms this notion by identifying lineage-specific genes, suggesting distinct ecological or evolutionary pressures on different lineages. The observed gene specificity in *P. stuartii* and Taxon 3, for example, hints at possible adaptive strategies unique to these taxa.

Conversely, the gene flow observed in Taxon 7 and *P. hangzhouensis* suggests potential interactions or shared evolutionary histories with other lineages. Furthermore, pan-genome analysis revealed noticeable core gene flow among different species, promoting the shaping of species diversity. The primary metabolic pathways and functional characteristics of the genus’s core genes were also discerned. The pathway associated with beta-lactam resistance could be related to the frequent acquisition of antibiotic-resistance genes within this genus, highlighting potential adaptive strategies in the face of antibiotic pressures.

The considerable presence of antibiotic resistance genes, coupled with their wide spectrum, highlights a potential frequent acquisition of such genes within species of this genus, typically ascribed to HGT events. Prior research has demonstrated the presence of multiple types of integron structures within the genus to carry resistance genes (17). In line with this, our study discovered that plasmids play a larger role in carrying resistance genes, with nearly half of these genes found to be plasmid-borne. Particularly, plasmid cluster 0 emerged as a significant locus, not only accounting for a remarkable proportion of the overall ARGs but also establishing itself as a primary reservoir for carbapenemase genes. This dominance suggests that cluster 0 may play a critical role in the dissemination of resistance genes, particularly carbapenem resistance genes, through plasmid-mediated transfer.

Upon delving into the species-specific resistance genes, we singled out three high-frequency resistance genes in *P. stuartii*, indicating they might be intrinsic to this species. Regarding the clinically significant carbapenem resistance genes, Taxon 3, *P. stuartii*, and *P. hangzhouensis* L12 displayed the greatest number of such genes along with diverse resistance genotypes. While this observation hints at a potential heightened clinical concern related to these species or lineages, it is important to note the limitations due to potential database biases and highlight the need for more comprehensive studies and continuous monitoring.

While our study provides valuable insights into the taxonomy and antibiotic resistance of the *Providencia* genus, certain limitations must be acknowledged. A substantial portion of our data was sourced from public databases. Although these repositories offer a wealth of genomic data, they can introduce biases. Specifically, strains exhibiting unique phenotypes, heightened resistances, or of particular clinical or scientific interest might be overrepresented due to targeted sequencing efforts. These selection biases can influence the observed distribution of resistance genes or specific genomic features. For instance, there is a possibility that resistant strains are sequenced more frequently, leading to an overrepresentation of resistance genes in publicly available datasets. Given these considerations, while our findings shed light on notable patterns and trends within the *Providencia* genus, the derived conclusions, especially regarding antibiotic resistance, should be interpreted with caution. Future studies aiming for a more holistic view of this genus would benefit from systematic, unbiased sampling, coupled with experimental validation and detailed epidemiological data.

In conclusion, this study addresses the previously perplexing taxonomical classification of species within the *Providencia* genus. It not only identifies a novel species—*Providencia zhijiangensis* sp. nov.—but also uncovers a synonymous pair, *P. stuartii* and *P. thailandensis,* in addition to expanding the genus number from 13 to 20. These findings significantly broaden the genomic landscape of *Providencia*, pointing out critical gaps in our understanding of the phylogenetic space it occupies. Through the investigation of core genes, we unveiled the primary metabolic pathways within the genus. Further explorations into antibiotic resistance genes and plasmids underscored the critical role of plasmids in carrying resistance genes. Additionally, the diversity of carbapenem resistance genes within three specific species or lineages signals the need for enhanced surveillance of these organisms in the future.

## Materials and methods

### Strain collection and whole-genome sequencing

To achieve a comprehensive evaluation of the *Providencia* genus, we adopted a holistic approach. Specifically, given the rarity of *Providencia* strains in our hospital, we obtained and sequenced four unique human-derived strains in 2022, representing the entirety of the samples available to us at that time. These strains were isolated from diverse clinical sources: two from urine, one from bile, and one from tissue fluid, reflecting a broad spectrum of clinical contexts.

Simultaneously, to ensure a comprehensive assessment of the *Providencia* genus, we extracted all available *Providencia* genomes from the NCBI’s GenBank database as of 1st December 2022, seeking to capture the broadest genomic diversity possible. This preliminary count indicated a total of 735 genomes. Recognizing the importance of data quality, we subjected each genome to a meticulous quality control assessment using QUAST v5.2.0 (40) and CheckM v1.2.2 (41). Genomes that failed to meet our quality standards (contig counts <300, N50 >50 kb, completeness >90%, contamination <2%) were excluded from subsequent analyses (**Supplementary Data 1**).

Phenotypic characterization of *Providencia zhijiangensis* sp. nov.

As described previously (42, 43), the Gram-stain was performed and the biochemical characteristics were determined using the bioMérieux API 20E and API 50CH kits according to the manufacturer’s instructions. Oxidase activity was determined using oxidase reagent (bioMérieux). The whole-cell fatty acids of strain D459 were assessed by the Guangdong Institute of Microbiology (Guangzhou, Guangdong, China). Meanwhile, in vitro antimicrobial susceptibility tests were conducted using Vitek II through broth microdilution. Breakpoints were determined using CLSI criteria (44), with the exception of tigecycline, for which the European Committee on Antimicrobial Susceptibility Testing (EUCAST; http://www.eucast.org/) guidelines were adopted.

### Genome assembly and OGRIs calculations

The genomic DNA of the isolates was extracted with the AxyPrep bacterial genomic DNA miniprep kit by Axygen Scientific, located in Union City, California, USA. The whole-genome sequencing was carried out using two platforms: the Illumina HiSeq 2500 system, which performed a paired-end run with 2 × 150 base pairs, and the Oxford Nanopore MinION platform. Raw sequencing reads underwent preprocessed with fastp v0.23.2 (45) for adapter sequences and low-quality bases trimming. Trimmed paired-end reads were subjected to assembly using shovill v1.1.0 (https://github.com/tseemann/shovill), with contigs shorter than 200 bp being filtered out.

The complete D4759 genome was assembled by integrating the short-read data from Illumina sequencing with long-read data from Oxford Nanopore sequencing using the Unicycler hybrid assembly pipeline (46). The 16S rRNA gene sequences of D4759 were obtained using PCR, with the application of universal primers 27F and 1492R (47). The related gene sequences from other species were sourced from the EzBioCloud database (26). All these 16S rRNA gene sequences were then aligned for comparison using MAFFT v7.505 (48). Subsequently, a phylogenetic tree was constructed using FastTree (49) based on the maximum likelihood method. Overall genome relatedness indexes (OGRIs) including ANI and in silico digital DNA–DNA hybridization index (dDDH) were assessed using the fastANI v1.33 (50) and genome-to-genome distance calculator (GGDC; formula 2) (24, 51), respectively. A cutoff of ≥96% ANI (23) or ≥70.0% in dDDH (23, 24) was employed to define a bacterial species. Further confirmation of the accurate bacterial species based on the results of TYGS database (52). Pairwise average amino acid identity (AAI) between genomes was computed using CompareM v0.1.2 (https://github.com/dparks1134/CompareM), with thresholds (65-72%) for genus-level ranks (28, 53).

### Pangenome analysis

All genomes were annotated with Bakta v1.7.0 (54). Pangenomes of all the isolates included in this study were analyzed using Panaroo v1.3.2 (55), with the annotated contigs in GFF3 format generated by Bakta as input. To generate the core-gene alignment, the parameters were set to a moderate model and a 95% threshold for protein sequence identity similarity and length difference cutoff as previously described (56). Additionally, MAFFT v7.505 (48) was specified with the ‘-a’ flag to align the core genes present in at least 95% of the genomes.

A pangenome gene classification, which is attuned to the population structure and based on within-species distribution, is derived using twilight package (29). Groups are delineated by the lineages established via fastbaps (as detailed below). The minimum size genome of the group is assigned a value of 1 to include all species.

### Phylogenetic analysis and population structure

A maximum likelihood (ML) phylogeny was constructed based on the core-genome alignment generated by Panaroo using RaxML v8.2.12 (57) with a GTR nucleotide substitution mode and 1,000 rapid bootstrap replications. The phylogeny was plotted and annotated using ggtree v3.7.1 (58). The core genome alignment and the phylogeny were subjected to fastbaps v1.0.8 (59) to infer population structure.

### Function classification

The COG functional categories of the gene sets for the *Providencia* genus were classified using eggNOG-mapper v2.1.9 (60). GO and KEGG ortholog assignment were annotated using InterProScan v5.56 (19) and KofamScan v1.3.0 (61), respectively.

### Identification of ARGs, and ISs

Antimicrobial resistance genes (ARGs) were identified using ABRicate v1.0.0 (https://github.com/tseemann/abricate) with a minimum identity threshold of 90% and a minimum coverage threshold of 90% based on NCBI AMRFinder Plus database. Insertion sequences (ISs) were annotated via ISfinder (62).

### Plasmid annotation and clustering

Plasmids from all assemblies were detected and reconstructed using the mob-recon tool from the MOB-suite toolkit v3.1.2 (63), employing default parameters. Plasmid clustering was undertaken using RabbitTClust v.2.2.1 (64) with a cutoff of a mash distance of 0.05.

### Data availability

The whole-genome sequences of four *Providencia* isolates from this study have been deposited in the GenBank database under BioProject accession PRJNA983084. The complete genome, including the chromosome and plasmids of strain D4759, has been submitted to NCBI GenBank under accession numbers CP135990-CP135992. All genome information used in this study can be found in **Supplementary Data 1**.

## Data Summary

The whole-genome sequences of four *Providencia* isolates from this study are available in the GenBank databases under BioProject accession PRJNA983084.

## Author contributions

XD and YZ designed the study. HQJ, YYY and YHX collected the isolates and clinical data. XD analyzed and interpreted the data. XD and HQJ wrote the manuscript. All authors reviewed, revised and approved the final manuscript.

## Acknowledgments

We would like to acknowledge all study participants and individuals who contributed to this study. We also acknowledge the support of National Infectious Disease Medical Center startup fund (Y.Z.) (B2022011-1), Jinan Microecological Biomedicine Shandong Laboratory project (JNL-2022050B), and Leading Innovative and Entrepreneur Team Introduction Program of Zhejiang (No. 2021R01012).

